# Untangling structural factors and evolutionary drivers in nascent polyploids

**DOI:** 10.1101/2020.12.21.423805

**Authors:** Julie Ferreira de Carvalho, Solenn Stoeckel, Frédérique Eber, Maryse Lodé-Taburel, Marie-Madeleine Gilet, Gwenn Trotoux, Jérôme Morice, Cyril Falentin, Anne-Marie Chèvre, Mathieu Rousseau-Gueutin

**Affiliations:** IGEPP, INRAE, Institut Agro, Univ Rennes, Le Rheu, France

**Keywords:** *Brassica napus* (oilseed rape), euploid selection, fertility, genome stability, homoeologous exchanges, meiotic behavior, polyploidy

## Abstract

1. Allopolyploids have globally higher fitness than their diploid progenitors however, by comparison, most resynthesized allopolyploids have poor fertility and highly unstable genome. Elucidating the evolutionary processes promoting genome stabilization and fertility is thus essential to comprehend allopolyploid success.
2. Using the *Brassica* model, we mimicked the speciation process of a nascent allopolyploid species by resynthesizing allotetraploid *B. napus* and systematically selecting for euploid individuals over eight generations in four independent allopolyploidization events with contrasted genetic backgrounds, cytoplasmic donors and polyploid formation type. We evaluated the evolution of meiotic behavior, fertility and identified rearrangements in S1 to S9 lineages, to explore the positive consequences of euploid selection on *B. napus* genome stability.
3. Recurrent selection of euploid plants for eight generations drastically reduced the percentage of aneuploid progenies as early as the fourth generation, concomitantly with a quasi disappearance of newly fixed homoeologous rearrangements. The consequences of homoeologous rearrangements on meiotic behavior and seed number strongly depended on the genetic background and cytoplasm donor.
4. The combined use of both self-fertilisation and outcrossing as well as recurrent euploid selection, allowed identification of genomic regions associated with fertility and meiotic behavior, providing complementary evidence to explain *B. napus* speciation success.

## INTRODUCTION

All living plants have experienced at least one episode of Whole-Genome Duplication (WGD) during their evolutionary history (Jiao *et al*., 2011; One Thousand Plant Transcriptome Initiative, 2019). This process tends to increase genetic and phenotypic diversity at various levels, and has been associated with greater fitness and more diverse ecological niches in polyploids compared to their diploid relatives (Selmecki *et al*., 2015; Baniaga *et al*., 2019). These observations are based on the successful outcomes of millions of years of evolution, however immediately after WGD, polyploids have to overcome several challenges due to bearing more than two sets of each chromosome. These sets of chromosomes can be more or less divergent depending on the occurrence of intra or interspecific hybridization before WGD leading to creation of auto or allopolyploids, respectively. To increase its chance of speciation, complex genome stabilization processes must occur rapidly after the formation of an allopolyploid species. First, strict bivalent formation is required for the formation of balanced gametes. However, in the case of allopolyploidy, redundant chromosomes coming from related parental species (homoeologous chromosomes) may pair with each other, and impact meiotic behaviour and plant fertility. Interestingly, the predominance of these events differs greatly among allopolyploids. Generally, multivalent associations are prevented during meiosis in most of natural allopolyploids (Grandont *et al*., 2013). To explain this mechanism, different hypotheses were proposed: either pre-existing genomic divergence between constitutive diploid progenitors impedes homoeologous pairing, or the presence of a genetic control in progenitors lead to a regulating mechanism of homoeolog pairing in the polyploid context (Mason & Wendel, 2020). For instance, genetic factors were shown involved in complete or partial homoeologous pairing control respectively in natural wheat allopolyploids and oilseed rape (for review Jenczewski & Alix, 2004; Griffith *et al*., 2006; Gonzalo *et al*., 2019). Contrastingly, in resynthesized allopolyploids (such as Brassica and Tragopogon), illegitimate pairing between homoeologous chromosomes is observed as soon as the first meiosis (Tate *et al*., 2006; Szadkowski *et al*., 2010). After numerous recombination events between homoeologs, their global genomic similarity tends to increase, leading to even more illegitimate COs as formulated by Gaeta & Pires (2010) under the term ‘polyploid ratchet’. As expected, because of the presence of univalents and multivalents in various germ cells, viable balanced gametes and fertility are extremely low in resynthesized allopolyploids exhibiting mispairing behaviour. Yet, the early structural dynamics linked to rearrangements between homoeologous genomes and their impact on meiotic instability and fertility stay unexplored in newly formed allopolyploids. Thus, deciphering the evolutionary processes that generated genome stabilization in natural allopolyploids and how this could be achieved in resynthesized ones is essential to fully comprehend early polyploid speciation processes.

*Brassica napus* L. (AACC, 2n=4x=38) is an allotetraploid resulting from the interspecific hybridization that took place ca. 7500 years ago (Chalhoub *et al*., 2014) between two closely related diploid species *B. rapa* (AA, 2*n*=2*x*=20) and *B. oleracea* (CC, 2*n*=2*x*=18). Spontaneous populations of *B. napus* were so far never uncovered. Recent studies have nonetheless identified the European origin of both A and C subgenomes of *B. napus* (Ha *et al*., 2019; Song *et al*., 2020). Although homoeolog pairing is limited in natural *B. napus*, different studies have repeatedly demonstrated the presence of numerous homoeologous translocations, limited in size but contributing to intraspecific diversity in *B. napus* (Chalhoub *et al*., 2014; Samans *et al*., 2017; Higgins *et al*., 2018; Song *et al*., 2020). Resynthesized crosses between the diploid Brassica species have been created to mimic the first steps of allopolyploid speciation to inform on the role of homoeologous rearrangements in meiosis control, genome stabilization and seed production. Homoeologous rearrangements and high levels of aneuploids have been observed in different resynthesized lines of *B. napus* (Song *et al*., 1995; Gaeta *et al*., 2007; Szadkowski *et al*., 2010; Xiong *et al*., 2011; Rousseau-Gueutin *et al*., 2017). Homoeologous rearrangements promote drastic genome instability as 50% of gametes may present homoeologous exchanges as soon as the first meiosis (Szadkowski *et al*., 2010). Depending on progenitors and type of gamete formation used in resynthesized crosses, homoeologous rearrangements amplify in a non-random fashion in the first generations (Szadkowski *et al*., 2011; Rousseau-Gueutin *et al*., 2017) and alter meiosis and seed production (Szadkowski *et al*., 2010, 2011; Girke *et al*., 2012; Jesske *et al*., 2013; Rousseau-Gueutin *et al*., 2017). Thus, it is paramount to investigate how the presence of non-reciprocal homoeologous exchanges impact meiosis and fertility: either by the size and position of these rearrangements along the genome, disturbing homoeologous pairing; or by modifying allele dosage and genetic mechanisms controlling meiosis and homoeologous pairing. Additionally, in the first generations, a significant proportion of aneuploid individuals were described causing supplementary instability (Xiong *et al*., 2011). These aneuploids result from chromosome mispairing and can alter gamete viability and consequently seed yield (Gaeta & Pires, 2010; Xiong *et al*., 2011). Selection of euploid individuals might thus, allow disentangling the consequences of aneuploidy from those of homoeolog rearrangements on meiotic behaviour and fertility.

So far, only rearrangements in the first generations after allopolyploidization have been studied in resynthesized *B. napus* without replicated lineages from the same S0 and without distinguishing aneuploids from euploid individuals. Here, we aim to unravel the fate of these genomic exchanges in selected euploid generations of different resynthesized *B. napus* allopolyploids. By eliminating all aneuploid individuals, we can disregard the effect of aneuploidy on meiosis and seed production to focus on the structural and functional impact of homoeologous rearrangements. The extent and consequences of these homoeologous rearrangements in *B. napus* were followed in four resynthesized lines resulting from different genetic backgrounds, reciprocal parental crosses and different mode of allopolyploid formation. For each cross, several independent S1 lines were created in order to follow the dynamics of fixed homoeologous rearrangements over eight generations propagated by single seed descent (SSD). We assessed meiotic behaviour and fertility and explored the role of fixed non-reciprocal homoeologous exchanges on these traits in 358 individuals. Overall, our results highlight that selection of euploid individuals led to the disappearance of newly fixed homoeologous rearrangements. We described homoeologous rearrangements having variable functional and structural impact on meiotic stability and seed production depending on the genetic background and cytoplasmic donor. We finally propose a model to explain genome stabilization process in natural *B. napus*.

## MATERIAL AND METHODS

### Production of resynthesized *B. napus* populations through repeated selection of euploid individuals

Resynthesized *B. napus* lines were created by crossing two different *B. oleracea* and *B. rapa* cultivars. For *B. oleracea* (2*n*=2*x*=18, CC), we used the doubled haploid lines *‘*RC34*’* (*B. oleracea* var. *alboglabra*) and ‘HDEM’ (*B. oleracea* var. *botrytis italica*). For *B. rapa* (2*n*=2*x*=20, AA), we used ‘Z1’ (*B. rapa* var. *trilocularis*) and ‘C1.3’ (belonging to a fodder variety named ‘chicon’ var. *rapifera*). A resynthesized *B. napus* named ‘RCC’ was created by first crossing the *B. oleracea* ‘RC34’ (mother plant) and *B. rapa* ‘C1.3’ (father plant) lines, resulting in the amphiploid ‘F1 hybrid RCC’ (2n=2x=19, AC) (Fig. 1). This F1 hybrid was subsequently somatically doubled using colchicine (Chèvre et al. 1989), leading to the resynthesized *B. napus* ‘RCC-S0’ (2n=4x=38). A reciprocal cross between the *B. rapa ‘*C1.3’ and the *B. oleracea* ‘RC34’ lines was also performed, leading to the amphiploid ‘F1 hybrid CRC’ and to the resynthesized *B. napus* ‘CRC-S0’ (after colchicine treatment). The two resynthesized ‘RCC’ and ‘CRC’ lines were selfed (hand pollination of floral buds before anthesis), producing the ‘RCC-S1’ and ‘CRC-S1’ progenies (Fig. 1). Thereafter 11 ‘RCC-S1’ and 10 ‘CRC-S1’ were selfed. These lineages were advanced by SSD until the eighth generation. At each generation, only one plant with 38 chromosomes (euploid) was randomly chosen from the set of euploid offspring. In some cases, lines did not produce any progeny, leading to the extinction of such lineage. In addition to ‘RCC’ and ‘CRC’, we produced resynthesized *B. napus* lines by firstly crossing the *B. oleracea* ‘HDEM’ line with the *B. rapa* ‘Z1’ line. From this cross, we obtained three amphiploid F1 hybrids (named ‘EMZ1’, ‘EMZ2’ and ‘EMZ3’) that were somatically doubled using colchicine, producing the EMZ1-S0’, ‘EMZ2-S0’ and ‘EMZ3-S0’ resynthesized lines. These three genetically identical EMZ lines were self-fertilized, producing the ‘EMZ1-S1’, ‘EMZ2-S1’ and ‘EMZ3-S1’ progenies. Then, five to eight plants from each lineage were advanced by SSD until S8 generation (Fig. 1). The resynthesized ‘UG EMZ’ *B. napus* populations were produced by crossing the F1 hybrid ‘EMZ1, 2 or 3’ with the corresponding resynthesized *B. napus* ‘EMZ1, 2 or 3 S0’, leading to the formation ‘UG EMZ1, 2 or 3 S0’ *B. napus* lines. Resulting polyploids are thus the result of a cross between a female unreduced gamete of the F1 hybrid ‘EMZ1’ and a male reduced gamete of ‘EMZ1-S0’ (Fig. 1). After selfing these lines, 4 to 15 S1 plants from each line were advanced by SSD to the S8 generation.

**Figure 1.**
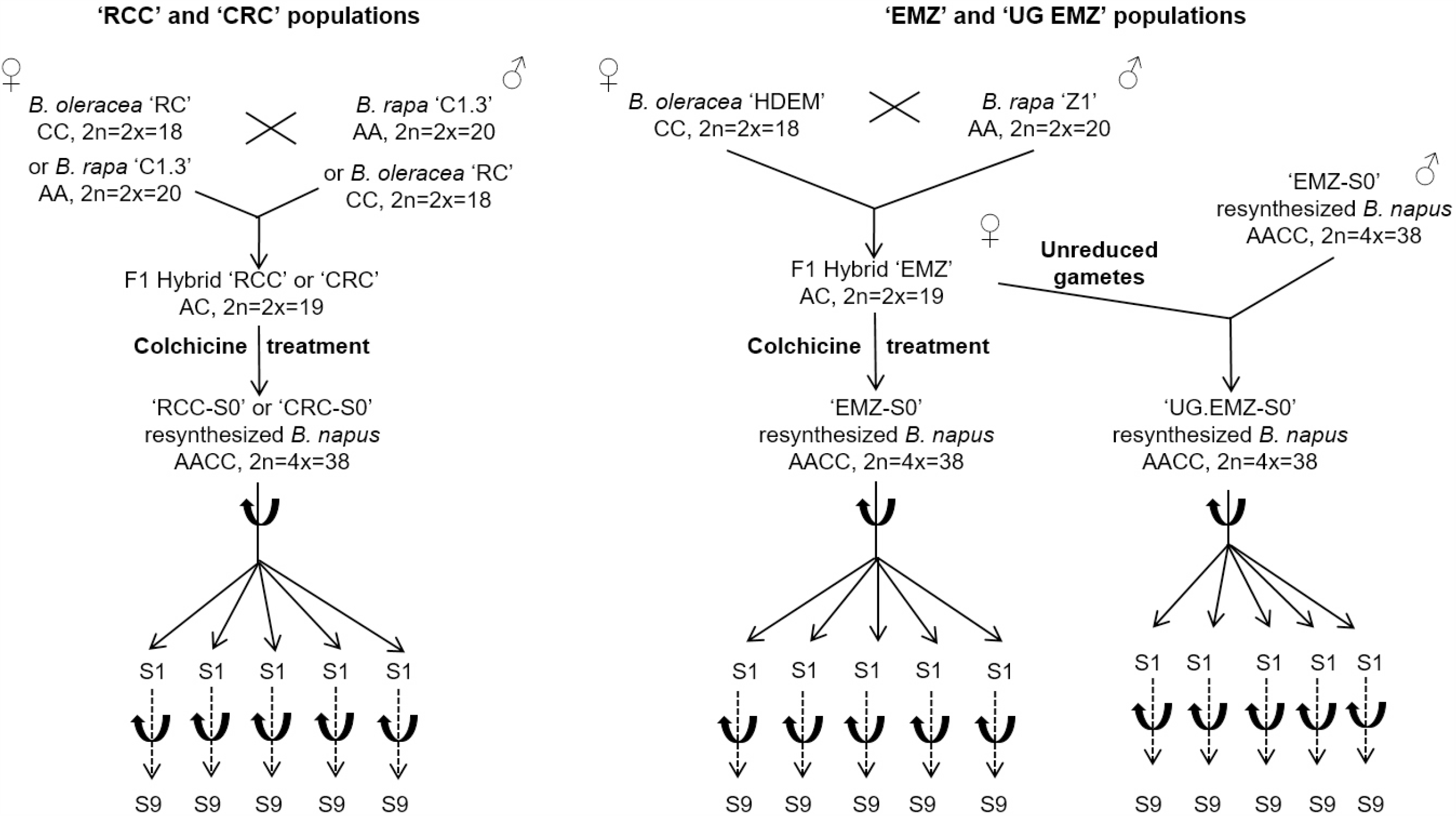
Crosses and gamete type formation of the different resynthesized *B. napus* populations advanced by single seed descent.

### Meiotic behavior and chromosome counting

The meiotic behavior and chromosome counting of all the produced material was studied by fixing floral buds in Carnoy’s solution (ethanol-chloroform-acid acetic, 6:3:1) for 24 hours and then stored in 50% ethanol. The anthers were squashed and stained in a drop of 1% acetocarmine solution: at least 20 pollen mother cells (PMCs) per plant were observed at metaphasis I.

### Fertility in resynthesized *B. napus* individuals

The fertility (number of seeds per 100 pollinated flowers) of the different resynthesized *B. napus* was calculated by counting the number of pods per pollinated flowers at the bud stage (preventing the impact of potential self-incompatibility in the parental lines, as notably already known for HDEM, Belser *et al*., 2018) and the number of seeds per pod allowing assessment of the number of seeds per pollinated flower. As a control, natural *B. napus* variety ‘Darmor’ was included in the experimental set-up.

### DNA extraction and SNP genotyping

Genomic DNA of 157 individuals was extracted with the maxi plant kit (LGC Genomics, Teddington Middlesex, UK) at the GENTYANE platform (INRAE, France). For all four resynthesized crosses, only generations S0, S1, S3, S6 and S8 were genotyped. Specifically, DNA was extracted from 42 individuals for the ‘RCC’ resynthesized population including the two diploid parents, the F1 hybrid, ‘RCC-S0’, and 38 resynthesized progenies (11 S1, 11 S3, nine S6 and seven S8). For ‘CRC’, DNA from 38 individuals was extracted including the F1 hybrid, ‘CRC-S0’, and 36 resynthesized progenies (ten S1, nine S3, nine S6 and eight S8). For the different ‘EMZ’ populations, 61 individuals were submitted to DNA extractions; the diploid parents, three F1 hybrids, three ‘EMZ-S0’, and 50 resynthesized progenies (18 S1, 16 S3, nine S6 and seven S8). Finally, DNA extractions on resynthesized lines created via female unreduced gamete pathway were performed on 16 individuals from ‘UG EMZ1-S0’ cross, 43 individuals from ‘UG EMZ2-S0’ cross and 16 individuals from the ‘UG EMZ3-S0’ cross.

Genotyping was then performed using the Illumina^®^ (http://www.illumina.com/) *Brassica* 60K Infinium SNP array (Clarke *et al*., 2016). Hybridizations were performed according to the standard procedures provided by the manufacturer. The obtained genotyping data were visualized using Genome Studio V2011.1 (Illumina, Inc., San Diego, CA, USA) and processed with a manually adapted cluster file.

### SNP data analyses

This *Brassica* 60K array is composed of 52,157 SNP markers that may either specifically hybridize to *B. rapa* or to *B. oleracea* or that may hybridize to both species. These latter markers thus hybridize to two distinct genome locations on *B. napus* (on the homoeologous A and C chromosomes). The positions of the SNP markers on the *B. napus* chromosomes derived from Rousseau-Gueutin *et al*., (2017) and were obtained by blasting the 52,157 sequence contexts (minimum of 90% global overlap and 90% identity) against the *B. napus* Darmor reference genome assembly (version 4.1 in Chalhoub *et al*., 2014). Only the markers presenting no more than one blast hit on each subgenome (A and C) were retained, enabling to discard SNPs potentially hybridizing at paralogous regions.

### Identification of deleted regions resulting from non-reciprocal homoeologous exchanges in each resynthesized *B. napus* line

To identify non-reciprocal homoeologous exchanges in the ‘RCC’ and ‘EMZ’ resynthesized populations, we used two types of markers: i) homoeo-SNP markers (Mason *et al*., 2017) that were homozygous and polymorphic (AA vs. BB) between the two diploid parental lines (*ie* ‘HDEM’ and ‘Z1’ or ‘RC’ and ‘C1.3’) and heterozygous in the S0 *B. napus* and ii) markers that only hybridized in one diploid parental line of the *B. napus* resynthesized population (i.e. “AA” in ‘HDEM’ versus ‘--’ in ‘Z1’). These markers specific to either *B. rapa* or *B. oleracea* are referred as dominant markers. Only the markers presenting identical genotype data for all four technical replicates were considered for further analyses. Thereafter, putative deletions in each resynthesized individual were identified by searching the loss of one parental allele in the polymorphic homoeo-SNP markers (‘AB’ in the S0 and ‘AA’ or ‘BB’ in the resynthesized *B. napus* progenies) or in the dominant markers (‘A-’ or ‘B-’ in the S0 and ‘--’ in the resynthesized *B. napus*). To avoid false positive results, deleted regions were only considered if at least three consecutive markers displayed the loss of one parental allele (from the same parent). Additionally, within a genealogy, deletions (with identical start and end positions, or extended start and/or end positions) had to be inherited from parents to offspring. This method allows for the detection of deleted regions from fixed non-reciprocal homoeologous DNA exchange in resynthesized *B. napus* polyploids (Rousseau-Gueutin *et al*., 2017). To perform these analyses, we developed a SQL database using pgAdminIII (v1.22.1) that was saved in postgresql v9.5 (DataBase Management System). The size and gene content of each deleted region was thereafter determined. We also evaluated whether the deletions were present in the distal region of a chromosome arm (last 30% of a chromosome arm) or close to the pericentromere (Mason *et al*., 2016).

### Statistical analyses

#### Multi-variable analyses and comparisons of means

In order to discriminate the different factors influencing both phenotyping (meiotic behavior and fertility) and genotyping variables (number, size and position of homoeologous rearrangements), we first conducted a Redundancy Analysis to summarize linear relationships between dependent variables and independent factors. Following this global analysis, statistical comparisons of means between crosses and between generations were performed using Anova and t-test with permutations when necessary. These statistical analyses and graphics were achieved using R language (R Core Team, 2017) and the RStudio environment (RStudio Team, 2015).

#### Probability distributions of typical measures of stability of meiosis and seed formation in newly resynthesized B. napus individuals

To statistically identify individual plants with extreme values of stability of meiosis and seed formation, we fitted probability density functions to the full cohort of observational data measured on the 358 resynthesized plants. The resulting probability distributions allowed weighting the probability of individual measure to conform or not to the expected pattern of meiosis and seed formation in newly resynthesized *B. napus*. Probability distributions were fitted using a classical approach of maximum likelihood estimate of parameters, by maximizing a log-likelihood function, with penalty applied for samples outside of range of the considered distribution. Considering the type and interval of measures, we fitted beta distributions for percentage measures (percentage of cells with 19 bivalents, with multivalents and percentage of male sterility), gamma distributions for mean positive measures (average number of univalent, bivalent and multivalent per PMC), binomial distributions for the ability to produce seeds and log-normal distribution for the number of seeds per 100 flowers.

#### Genome scan for linking extreme phenotypic measures and deleted sites

To identify putative genomic areas implied in the stability of meiosis and seed formation, we scanned the genome for deleted SNPs included in structural rearrangements (i.e deleted regions from one subgenome that most presumably result from homoeologous exchanges). We hypothesized that plants measured with extreme phenotypes may present specific deleted genomic areas involved in the stability of meiosis and seed formation. We thus classified for each SNP position, all studied plants in two groups: plants with or without this particular deleted SNP, and analyzed their phenotypic measures. We compared phenotypic measures of plant with and without deletions, and then computed the probability of overrepresentation of each identified deletion in plants with extreme phenotypic values.

In the first step, we tested if the two groups of plants, with and without deletion, differed for their phenotypic measures using classical Mann-Whitney rank tests. The advantage of such non-parametric test is to be less likely to find false significant differences than using parametric test, and thus identifying robust candidate regions. When such identified deletions occurred in different lineages with different genetic backgrounds, it strengthened our confidence that such deletion may include candidate genes involved in the stability of meiosis or seed formation. Indeed, different genetic backgrounds with the same deletion and the same phenotypic measures decrease the probability that identified extreme measures could be due to another deleted site co-inherited by descent from a common ancestor. We thus performed two complementary variations of this approach: one *overall data* and one *within each genetic cross*. First, we review each deletion regardless of their genetic background and tested if plants with deletion conformed to the standard distribution of phenotypic measures or not. Second, in each of the genetic combination, we tested if plants with deletion conformed to the standard distribution of phenotypic measures or not for each deletion, allowing to identify cross-specific loci that may be co-localized.

In the second step, we identified plants with extreme phenotypic measures when their measures lied outside the 99% confidence interval of each phenotypic measure, considering the appropriate fitted probability distributions (see above). Then, for each genomic site *i*, we counted the number of time *k*_*i*_ this one was deleted among the *n*_*i*_ abnormal plants. We computed the probability to observe *k*_*i*_ deleted sites among *n*_*i*_ by chance *P*(*k*_*i*_/*n*_*i*_, *p*_*i*_) as the probability mass function of a binomial distribution of success probability in each trial as the overall ratio of deleted sites on the number of successfully genotyped sites. A SNP candidate was considered as involved in the phenotype if its probability *P*(*k*_*i*_/*n*_*i*_, *p*_*i*_) was inferior to 10% in at least six individuals in different genetic backgrounds.

## RESULTS

The dataset presented in this study comprises phenotypic measures of SDD individuals over eight generations for four independent nascent lineages of allopolyploid *B. napus*. The impact of repetitive euploid selection was visible as soon as the fourth generation and forward, as only 0 to 2.94% of aneuploid individuals were found in the resynthesized S4 to S8 generations compared to 11.46-11.27% in S1-S3 (Supporting Information Fig. S1). In total, we assessed the meiotic behavior and seed yield of 358 individuals including 73 individuals for each ‘RCC’ and CRC’ lines, as well as 91 and 121 ‘EMZ’ and UG EMZ’ individuals, respectively. All individual measurements included number of seeds per 100 flowers, percentages of cells with multivalents and bivalents as well as, mortality in the next generation (Supporting Information Table S1). This valuable dataset was then analysed in regards to a genotyping dataset performed on the four lines at generation S1, S3, S6 and S8. Each resynthesized *B. napus* lineage is represented by 35, 37, 40 and 62 ‘RCC’, ‘CRC’, ‘EMZ’ and ‘UG EMZ’ individuals, respectively. Description (number, size, position) of the identified homoeologous rearrangements was included in Supporting Information Table S1. Overall, variability of the dataset was well explained (68%, p<0.001) by the factors ‘cross’ and ‘generation’, as well as by the interaction of ‘cross’ and ‘generation’ (p<0.001; Supporting Information Table S2). We thus mined the datasets to identify the factors explaining the phenotypic and genomic variations between genetic backgrounds as well as the dynamics in the first eight generations after allopolyploid speciation.

### Fertility

The fertility of the resynthesized *B. napus* populations was assessed based on the number of seeds per 100 flowers in all four nascent lines of *B. napus* ‘RCC’, ‘CRC’, ‘EMZ’ and ‘UG EMZ’. Means for the nascent lines were 270.8, 158.8, 62.6 and 55.1, respectively. By comparison, variety ‘Darmor’ has on average 2067 seeds per 100 flowers (SD=516) which is significantly 10-fold higher than what is observed in resynthesized *B. napus* (t-test with permutation, p=0.0025, Supporting Information Fig. S2). Overall, differences in seed yield were significant between all lines (t-test, p<0.01) except between ‘EMZ’ vs ‘UG EMZ’ (Supporting Information Fig. S2). In ‘RCC’, fertility significantly decreased from the 1^st^ generation to the 4^th^ (t-test, p<0.01) and again from the 5^th^ to the 8^th^ generation (t-test, p<0.01) with a high fertility (mean=572.2) observed in the 5^th^ generation (comparable to the yield observed in S1) (Fig. 2). In ‘CRC’, fertility increased in the 2nd generation compared to the first generation (162.5 to 395.8 seeds per flower, p<0.05) to drastically decrease until the 8^th^ generation (t-test, p<0.05) (Fig. 2). In ‘EMZ’, compared to S0 all following generations presented a lower number of seeds. Similarly, ‘UG EMZ’ individuals exhibited equal or lower number of seeds in the generations following allopolyploidization, only a slight significant decrease was observable from generation S1-S2 to S3 (t-test, p<0.05) (Fig. 2).

**Figure 2.**
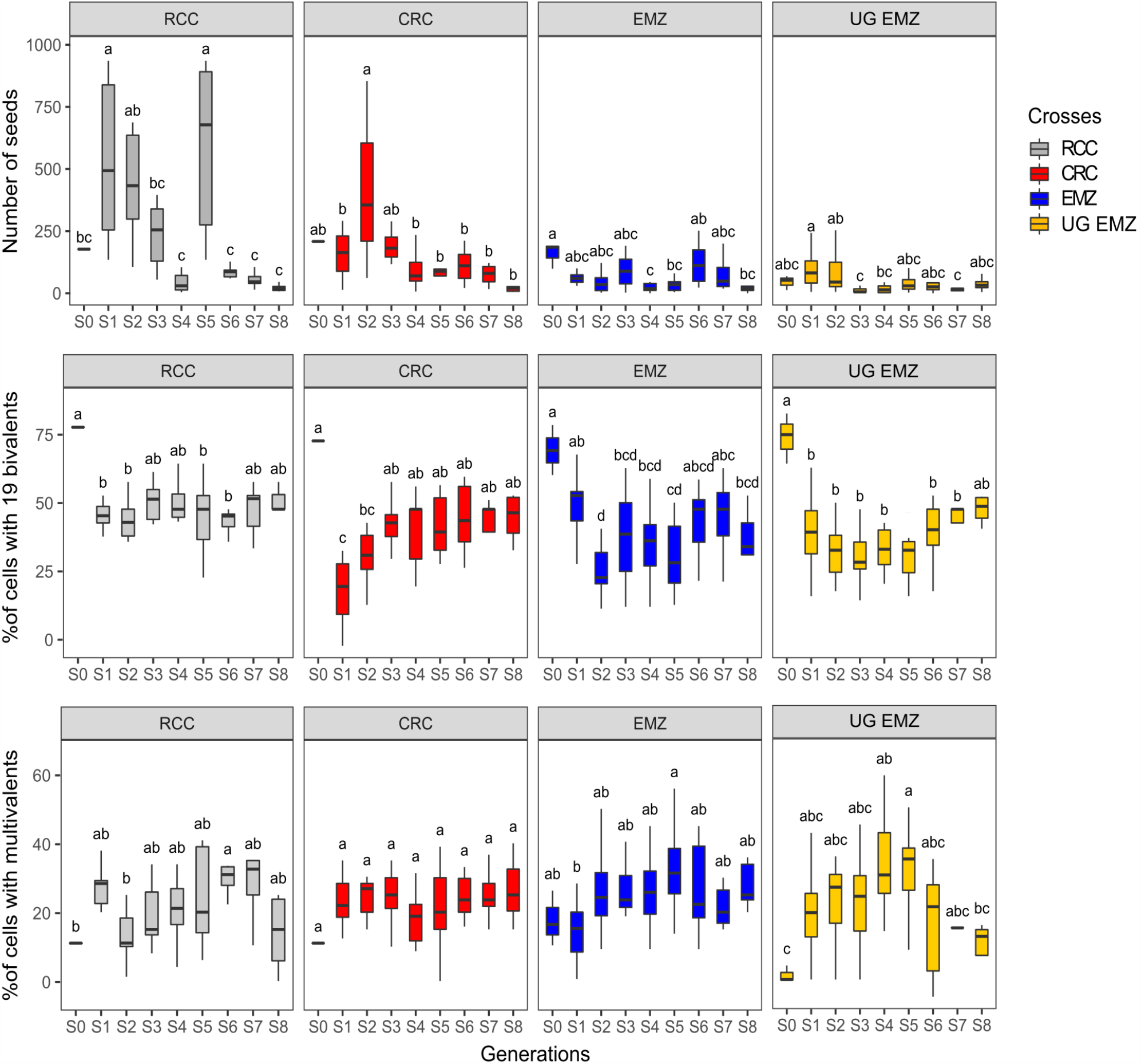
Evolution of number of seeds per 100 flowers, percentage of cells with 19 bivalents and percentage of cells with multivalents during Metaphasis I in all four resynthesized populations across all generations. Letters above boxplots represent significativity after t-test (p<0.05).

### Meiotic behavior

Meiotic behavior was characterized using various descriptors. First, as solely euploid *B. napus* individuals (with 2n=38) were assessed, it was expected that such plants presented a better meiotic stability compared to its aneuploid siblings. Despite having 38 chromosomes, meiosis was affected in our plants with 49.4%, 41.0%, 41.1 and 38.1% of cells exhibiting 19 bivalent structures in ‘RCC’, ‘CRC’, ‘EMZ’ and ‘UG EMZ’, respectively overall generations (Supporting Information Fig. S2). Percentage of cells with 19 bivalents was significantly higher in ‘RCC’ compared to the three other nascent lines (Supporting Information Fig. S2, t-test, p<0.01).

S0 individuals in all nascent lines exhibited overall higher percentages of cells with proper bivalents (80.0% in RCC, 75.0% in CRC, 71.6% in EMZ and 76.3% in UG EMZ) compared to subsequent generations. Percentage of cells with 19 bivalents was consistent across generations S1 to S8 in ‘RCC’ whereas significant differences between generations were visible in the crosses ‘CRC’, ‘EMZ’ and ‘UG EMZ’ (Fig. 2). In ‘CRC’, percentage of cells with 19 bivalents tended to increase in S3 compared to S1 (from 21.1% to 44.8%, t-test, p<0.05, Fig. 2) and stabilized in subsequent generations. In ‘EMZ’, percentage of cells with 19 bivalents decreased drastically from S1 to S2 (t-test, p<0.05, Fig. 2) followed by a slight increase from S2 to S7. In ‘UG EMZ’, after the drop from S0 to S1, percentage of cells with 19 bivalents was found constant with a slight increase in S8 (Fig. 2).

Finally, we assessed the percentage of cells exhibiting multivalents in the nascent allopolyploids. A similar percentage of cells with multivalents was observed among all lines (27.3% in ‘RCC’, 28.8% in ‘CRC’, 29.7% in ‘EMZ’ and 29.1% in ‘UG EMZ’, Supporting Information Fig. S2). This trend was also found consistent across generations for all resynthesized lines except for ‘UG EMZ’ where a slight increase in the percentage of multivalents was observed from S1 to S4-S5 (23.4% to 34.7%, t-test, p<0.05, Fig. 2). After that, the predominance of multivalents decreased from S5 to S8.

### Identification of fixed non reciprocal structural rearrangements

To identify the putative presence of fixed homoeologous rearrangements in each genotype, we used the polymorph markers between the diploid parental lines of resynthesized *B. napus*. For the ‘RCC’ and ‘CRC’ populations, a total of 12,218 markers including 2,274 co-dominant and 9,944 dominant markers were included in the analysis (one marker every 68.8kb along the assembled *B. napus* genome). Similarly, 15,180 markers were analysed in the ‘EMZ’ and ‘UG EMZ’, including 13,501 dominant markers and 1,679 co-dominant markers (one marker every 55.3kb along the assembled *B. napus* genome).

We then evaluated size, number and position on the chromosomes and on the subgenomes of fixed homoeologous rearrangements in a population per resynthesized line and per generation (Supporting Information Table S1), using the physical localization of SNP markers on *B. napus* genome (Rousseau-Gueutin *et al*., 2017). Average number of identified regions ranged between 5.2, 6.1, 8.1 and 11.2 per individual per generation in ‘RCC’, in ‘CRC’, in ‘EMZ’ and ‘UG EMZ’ respectively (Fig. 3). Globally, individuals from the ‘UG EMZ’ line showed significantly higher number of rearranged regions than ‘CRC’ and ‘RCC’ lines (t-test, p<0.01, Fig. 3). The average size of the homoeologous rearrangements was estimated at 1.88, 3.79, 2.43 and 3.06Mb in ‘RCC’, in ‘CRC’, in ‘EMZ’ and ‘UG EMZ’ respectively (Fig. 3). A significant difference was only observed between ‘RCC’ and ‘UG EMZ’ (t-test, p<0.01, Fig. 3). We observed a limited number of homoeologous rearrangements in ‘RCC’ and ‘CRC’ compared to other crosses, with the exception of one individual in ‘CRC’ having a larger number of rearrangements in S6 and S8 (Fig. 3). By contrast, number of homoeologous rearrangements is higher as soon as the S1 generation in ‘UG EMZ’ and to a lesser extent in ‘EMZ’. We observed a low number of individuals in generations S8 in both ‘EMZ’ and ‘UG EMZ’ compared to ‘RCC’ and ‘CRC’. Interestingly, by decomposing the nature of these homoeologous rearrangements for each individual compared to previous generation, we could infer the drastic decrease of new homoeologous rearrangements appearing in the different lineages at S3 for ‘RCC’ and at S6 for ‘CRC’, ‘EMZ’ and ‘UG EMZ’ (Supporting Information Fig. S3).

**Figure 3.**
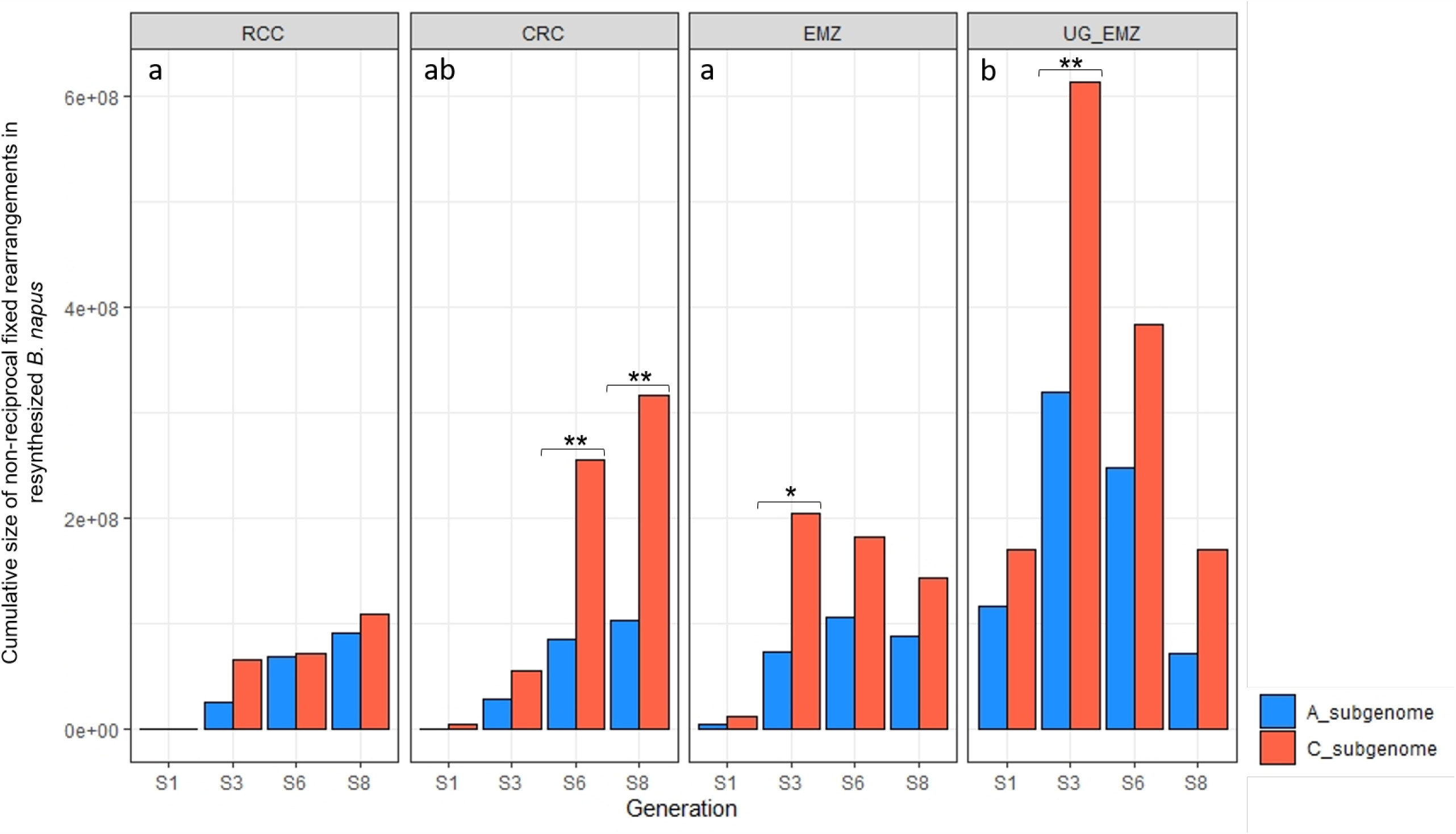
Structural dynamics of non-reciprocal homoeologous rearrangements (a) at each generation for all four resynthesized *B. napus* populations (lines connect data point from the same lineage). Per population, we also compared (b) the number of homoeologous rearrangements and (c) the average size of these rearrangements across all individuals. Letters above boxplots represent significativity after t-test (p<0.01).

In parallel, we assessed cumulative size of homoeologous rearrangements on each subgenome from generation S1 to S8 in all crosses (Fig. 4). Overall, cumulative size of homoeologous rearrangements was found higher in ‘UG EMZ’ compared to ‘EMZ’ and ‘RCC’ (with 35.2Mb vs 20.5Mb and 12.4Mb; t-test, p<0.05) (Fig. 4). Furthermore, the C subgenome was predominantly affected by rearrangements in ‘CRC’, ‘EMZ’ and ‘UG EMZ’ (Fig. 4) but only statistically different in generations S6 and S8 of ‘CRC’, generation S3 of ‘EMZ’ and S3 of ‘UG EMZ’. Although having similar progenitors, crosses ‘RCC’ and CRC’ were found differently impacted by homoeologous rearrangements on the C subgenomes in S6 and S8 (t-test, p<0.05).

**Figure 4.**
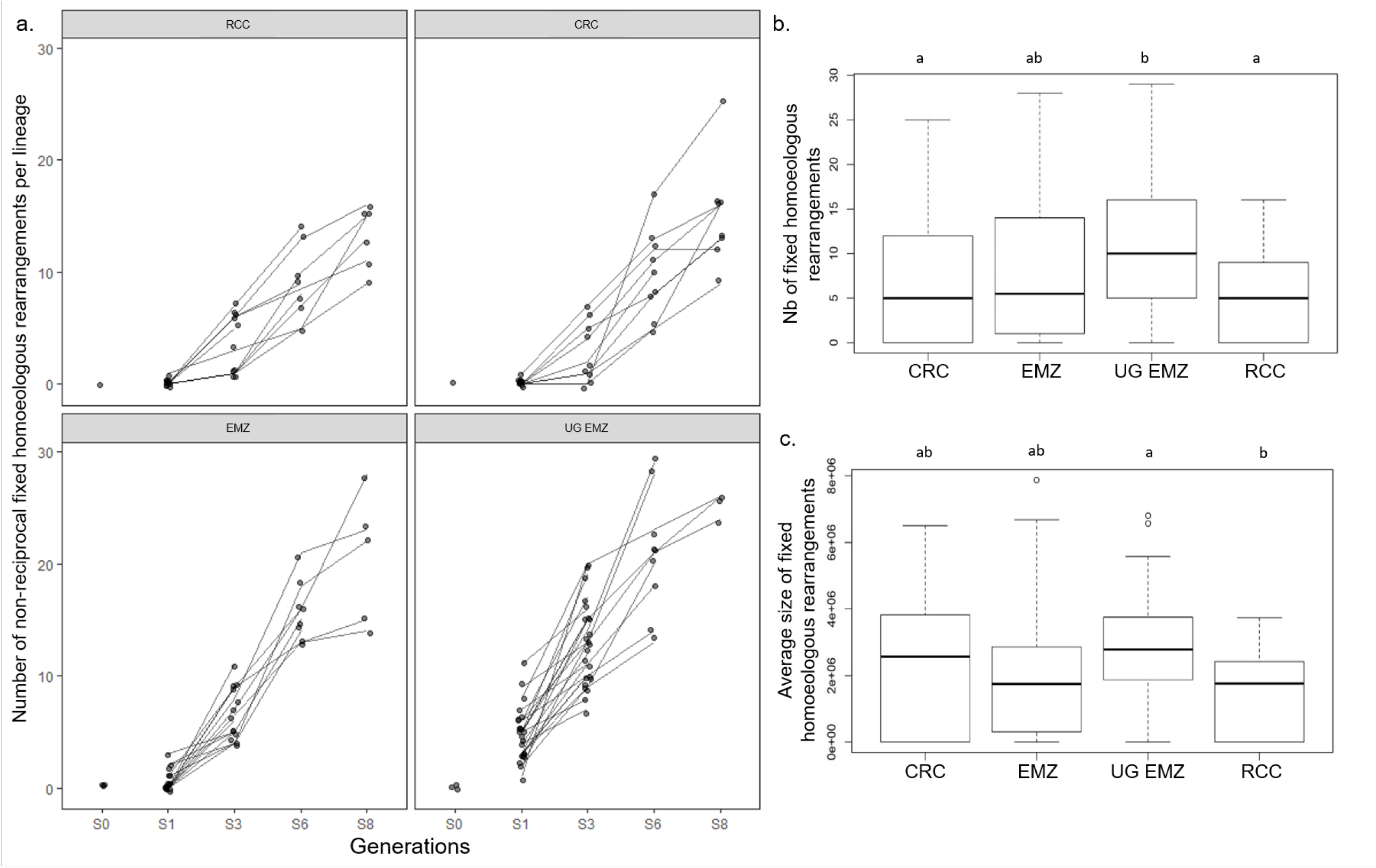
Cumulative size of translocations in A and C subgenomes in all individuals at each generation for all four resynthesized *B. napus* populations. Letters in each graph represent the overall significant differences between populations (t-test, p<0.05) whereas the stars represent significant differences in cumulative size between subgenomes in a generation (t-test, p<0.05).

### Correlations between homoeologous rearrangements, meiotic behaviour and fertility

The novelty of the present study lies in explaining meiotic behavior, fertility, and structural dynamics using repeated euploid selection of *B. napus* individuals in nascent allopolyploid populations. We assessed correlations between the different variables in order to identify if and how fixed non-reciprocal translocations may explain seed yield and chromosome pairing during meiosis. First, all measurements describing the presence of homoeologous rearrangements were correlating positively between themselves (Supporting Information Fig. S4). Expectedly, percentage of cells with bivalents was inversely correlated with percentage of cells with multivalents (with statistical support in crosses ‘RCC’, ‘EMZ’ and ‘UG EMZ’, Supporting Information Fig. S4). Interestingly, the correlations depended profoundly on the cross and thus the progenitors of the resynthesized allopolyploids. In ‘RCC’ and ‘CRC’ that have globally fewer homoeologous rearrangements, strongest negative correlations were found between the presence of homoeologous rearrangements and the number of seeds produced (Supporting Information Fig. S4). By contrast, large and numerous rearrangements observed in ‘EMZ’ have strong negative impact on meiotic behavior (Supporting Information Fig. S4). In ‘UG EMZ’, correlations were described between average size of rearrangements and meiotic behavior and between number of homoeologous rearrangements along with cumulative size on A subgenome and fertility (Supporting Information Fig. S4).

### Genome scan linking phenotypic measures with rearranged homoeologous regions

To go further, a genome scan was performed to link phenotypic measures with positions of the rearranged homoeologous regions (Supporting Information Table S2). Using genome scan analysis *overall data*, we identified one locus of 2.2Mb on chromosome A03 containing 440 genes (Table 1) and impacting the percentage of cells with multivalents. This list of genes was screened to identify annotations that might be relevant in the context of plant meiosis. In particular, one gene (ortholog AT2G42890) coding for a protein MEI2-like 2 or ML2 has drawn our attention. This gene has several copies in the genome of *B. napus* ‘Darmor’ (with two paleologs BnaA03g19940, BnaC03g23940/BnaC03g23930 and BnaA05g03150, BnaCnng35790). The unique gene on chromosome A03 has potentially been replaced by the two copies from chromosome C03. Using genome scan analyses *within each cross*, we identified one region associated with the percentage of cells with 19 bivalents in ‘EMZ’ and ‘CRC’ individuals on the C02 chromosome and three regions associated with the number of seeds per 100 flowers on C01, C02 and C04 chromosomes (containing 228 to 880 genes, Table 1).

**Table 1.** Summary of homoeologous deletions having a significant impact on percentage of cells with univalents/multivalents and on the number of seeds per 100 flowers in resynthesized *B. napus* individuals (see statistical analyses part). The physical localization of the identified candidate region as well as the number of SNP markers and number of genes in the region are indicated. We also mentioned for each population the number of individuals harboring the deletion of the candidate region via a homoeologous rearrangement, average phenotypic measures and standard deviation in brackets. Genes implicated in meiosis were retrieved using meiosis gene list from Higgins et al. (2020). Significant impact on phenotypes is represented by (*) when the deletion was significantly associated with higher or lower phenotypic measures in one or several genetic backgrounds.

| Phenotypic trait | Chromosome | Start position | End position | Number of SNPs | Number of genes | Genes implicated in meiosis | 'EMZ' individuals |  | 'UG EMZ' individuals |  | 'RCC' individuals |  | 'CRC' individuals |  | All individuals |  |
| --- | --- | --- | --- | --- | --- | --- | --- | --- | --- | --- | --- | --- | --- | --- | --- | --- |
|  |  |  |  |  |  |  | With deletion | Without deletion | With deletion | Without deletion | With deletion | Without deletion | With deletion | Without deletion | With deletion | Without deletion |
| % of cells with multivalent | A03 | 7,593,332 | 9,793,914 | 16 | 440 | 1 (ML2, AT2G42890) | 2 | 40 | 3 | 58 | 4 | 27 | 2 | 32 | 11 (*) | 157 |
|  |  |  |  |  |  |  | 0.18%(0.6%) | 0.26%(0.11%) | 0.12%(0.16%) | 0.26%(0.14%) | 0.22%(0.16%) | 0.29%(0.09%) | 0.24%(0.06%) | 0.30%(0.09%) | 0.19%(0.14%) | 0.27%(0.11%) |
| % of cells with 19 bivalents | A02 | 17,970,614 | 20,069,824 | 23 | 258 | 0 | 5(*) | 36 | 6 | 50 | 1 | 30 | 8(*) | 26 | 20 | 142 |
|  |  |  |  |  |  |  | 0.23%(0.14%) | 0.51%(0.14%) | 0.38%(0.10%) | 0.40%(0.16%) | 0.50% | 0.50%(0.09%) | 0.51%(0.08%) | 0.37%(0.16%) | 0.40%(0.15%) | 0.44%(0.16%) |
| Number of seeds/100 flowers | C01 | 2,475,667 | 8,275,217 | 21 | 820 | 1 (CDC20,1, AT4G33270) | 5(*) | 40 | 10(*) | 52 | 4 | 30 | 8(*) | 29 | 27 | 151 |
|  |  |  |  |  |  |  | 53.69(57.54) | 90.73(62.16) | 44.60(49.66) | 60.90(76.16) | 108.53(117.02) | 266.00(268.61) | 60.68(69.70) | 161.26(135.93) | 55.70(63.42) | 135.44(168.94) |
| Number of seeds/100 flowers | C02 | 4,887,388 | 6,416,485 | 17 | 233 | 0 | 7 | 38 | 12(*) | 50 | 8(*) | 26 | 5(*) | 32 | 32 | 146 |
|  |  |  |  |  |  |  | 97.26(68.02) | 80.31(61.76) | 111.57(108.51) | 47.06(57.23) | 23.24(5.53) | 269.17(261.75) | 24.99(3.12) | 149.62(131.65) | 82.51(82.77) | 121.43(160.71) |
| Number of seeds/100 flowers | C04 | 44,901,241 | 46,386,410 | 45 | 228 | 0 | 9(*) | 36 | 12(*) | 50 | 2 | 32 | 7(*) | 30 | 30 | 148 |
|  |  |  |  |  |  |  | 43.00(44.28) | 92.04(62.82) | 16.11(22.03) | 63.85(72.05) | 181.01(162.39) | 251.63(265.50) | 62.33(59.33) | 157.52(136.74) | 47.13(65.05) | 129.60(161.82) |

## DISCUSSION

In this study, we unravel the consequences of repeated euploid selection in the early generations following allopolyploidization, specifically we describe how rearrangements between homoeologous chromosomes influence meiotic behavior and fertility in nascent resynthesized *B. napus* individuals. Compared to previous studies using resynthesized *B. napus* (Gaeta & Pires, 2007; Szadkowski *et al*., 2010; Xiong & Pires, 2011; Xiong *et al*., 2011; Rousseau-Gueutin *et al*., 2017; Bird *et al*., 2020), we decided to perform repetitive selection for euploid individuals in order to evaluate the role of this phenomenon in *B. napus* speciation success. Ultimately, after a few euploid generations the consequences on meiosis and fertility can be attributed primarily to homoeologous rearrangements thus drawing conclusion on the impact of homoeologous rearrangements on genome stabilization. After the first meiosis and its ‘genome blender’, we focus on the drastic subsequent increase of homoeologous rearrangements from S1 to S4, rapidly followed by a quasi total disappearance of newly fixed homoeologous rearrangements in all studied resynthesized *B. napus* lineages. Interestingly, their consequences on meiotic behavior and seed number strongly depended on the genetic background and cytoplasm donor. Finally, we discuss the origin of *B. napus* and the processes that could explain their genomic stability compared to resynthesized allopolyploids.

### 1. Impact of the first meiosis depends on the different genetic backgrounds

Immediately after the formation of the different resynthesized *B. napus* lines, S0 individuals exhibited a relatively high percentage of cells with bivalents and low percentage of cells with multivalents due to the absence of rearrangements (Szadkowski *et al*., 2010, 2011). In the S1 generation, percentage of cells with 19 bivalents significantly decreased while homoeologous rearrangements were fixed, favoring subsequent homoeologous pairing notably via the formation of multivalents.

The extent of homoeologous rearrangement intensification in the first couple of generations after allopolyploid speciation depends on the genetic background. Number of homoeologous rearrangements and cumulative size in S3 is strongly limited in ‘RCC’ and ‘CRC’ compared to ‘EMZ’. These differences can be explained by the contrasted genetic backgrounds of resynthesized *B. napus* progenitors. First, macrosyntenic differences (large structural variants) between progenitor genomes could be present leading to different rates of homoeologous pairing and crossing-over in the allopolyploid. Both *Brassica rapa* used in this study (‘Z1’ ssp trilocularis and ‘C1.3’ ssp rapifera) and both *B. oleracea* progenitors (‘HDEM’ ssp botrytis ‘broccoli’ type vs ‘RC34’ ssp alboglabra ‘chinese kale’) come from distinct clades (Cheng *et al*., 2016). Although, the genome structure of ‘C1.3’ and ‘RC34’ is still unknown, a previous RNA-Seq study on the same resynthesized polyploids demonstrated the divergence in terms of transcription between progenitors and different transcriptional dynamics in the allopolyploids (Ferreira de Carvalho *et al*., 2019). Globally, large structural diversity is being described within both *B. rapa* and *B. oleracea* species (Lin *et al*., 2014; Golicz *et al*., 2016; Boutte *et al*., 2020) and could participate in the variation in number and size of homoeologous rearrangements. Second, presence of a genetic factor controlling meiotic behavior in allopolyploids and preventing homoeologous pairing could also explain the differences in the number of rearrangements observed and in the percentage of cells with bivalents between ‘RCC’ and ‘EMZ’. One approach consisted in correlating the loss of one homoeologous region with the variation observed in the different phenotypic variables. We notably identified one region on the A03 chromosome containing 440 genes in ‘RCC’, ‘CRC’ and ‘UG EMZ’ and leading to fewer cells with multivalents. Interestingly, one meiotic gene is included in this region and has been shown implicated in meiosis and control of chromosome pairing. *ML2* gene has been shown to play a role in fertility and plant meiotic behavior in *A. thaliana* mutants (Kaur *et al*., 2006). In RNAi triple mutants for *aml1, aml2* and *aml4*, phenotypes observed were due to a range of abnormalities in chromosome organization during meiotic prophase and later stages. Here, resynthesized lines of *B. napus* were not subjected to a complete loss of the locus but the replacement of the A03 region by the homoeologous region from the C subgenome. As the synteny is well-conserved between homoeologous regions, we can speculate that the homoeologous C allele is duplicated. Thus, as the function is conserved, allelic or transcription variation can be at the origin of the contrasted phenotypes. Interestingly, Kaur and co-authors, also suggested a dosage effect of the *ML1* to *ML5* genes as they reported a strong expression of these genes in meiocytes compared to other developmental stages and organs. Similarly, other genes have been described as implicated in homoeologous pairing control *via* dosage or allele effect, such as *MSH4* (Gonzalo *et al*., 2019). *BnaPH1* and *PrBn* have additionally been shown to reduce homoeologous pairing (Jenczewski *et al*., 2003; Higgins *et al*., 2020).

Finally, with identical diploid progenitors, resynthesized individuals ‘EMZ’ and ‘UG EMZ’ exhibited similar fertility and meiotic behavior but showed contrasted patterns regarding their respective cumulative size of fixed homoeologous rearrangements. These two resynthesized allopolyploids differ only by their mode of polyploid formation. The S0 from ‘EMZ’ has been produced *via* colchicine treatment whereas ‘UG EMZ’ S0 has been produced by unreduced gametes following the hybridization of one ‘EMZ’ S0 and one ‘EMZ’ F1 hybrid. Unreduced gametes formed in allohaploids (n=2x=19; AC) present an elevated frequency of homoeologous exchanges during meiosis compared to allotetraploids *B. napus* (2n=4X=38; AACC) (Nicolas *et al*., 2007, 2009, 2012; Cifuentes *et al*., 2010; Szadkowski *et al*., 2011). Indeed, ‘UG EMZ’ S3 individuals, cumulating rearrangements generated from unreduced female gametes with the ones occurring in S0 synthetic male gamete, have two to three-fold higher number of rearrangements and larger cumulative size than ‘EMZ’ S3 individuals. However, this tendency diminishes in S6 and S8 in ‘UG EMZ’ individuals thanks to the low fixation of novel homoeologous rearrangements over generation via the recurrent selection of euploid individuals as well as the high mortality rate of individuals with numerous rearrangements.

### 2. Consequences of euploid selection on the drastic decreased fixation of homoeologous rearrangements in early allopolyploid generations

Rapidly after the third generation, euploid selection led to the fixation of fewer novel homoeologous rearrangements hence, limiting the negative impact of these rearrangements on genomes stability and seed yield. Indeed, euploid selection seems to foster higher or at least constant number of bivalents during meiosis without observable consequences on seed yield. However, the amplitude of this result highly depends on the genetic background and maternal progenitor of the resynthesized allopolyploids.

Rearrangements between subgenome impact meiosis, the smaller and fewer the rearrangements in a plant, the more stable its meiosis. ‘RCC’ had fewer, shorter rearrangements without improved seed yield in S8, which probably reveals the functional impact of homoeologous rearrangements on this trait. As numerous seed yield QTLs were identified along the genome of *B. napus* (Raboanatahiry *et al*., 2018), we can presume that even few rearrangements in those regions will have an impact on the production of seeds. In this case, the decrease in seed number may be linked to the allelic diversity and/or dosage effect of genes present in the homoeologous rearrangements. This conclusion is also supported by the fact that in generations S1, S2, S3 and S5 some individuals exhibited a high number of seeds which indicate that some of the rearrangements present in those individuals have a beneficial or at least, a less deleterious impact than the others on seed yield. On the other hand, ‘CRC’ demonstrates fewer and shorter rearrangements with poorer seed yield than ‘RCC’. With identical genetic backgrounds, both crosses should be similarly functionally impacted by homoeologous rearrangements on seed production. Interestingly, ‘CRC’ individuals have additional homoeologous rearrangements on the C subgenome in generation S6 and S8 compared to ‘RCC’. This observable difference between ‘RCC’ and ‘CRC is probably due to the maternal donors. Phenotypic consequences of differential cytoplasm donors have been reported (in Brassica Cui *et al*., 2012 and in tomato Demondes de Alencar *et al*., 2020) and recently in maternal cross combination of resynthesized *B. napus* (Sosnowska *et al*., 2020). However, the underlying genetic interactions between organelle and nuclear genomes, as well as their impact on plant functional traits are still overlooked.

In both ‘EMZ’ and ‘UG EMZ’, large and numerous homoeologous rearrangements were observed, directly influencing the stability of meiosis and decreasing the percentage of cells with 19 bivalents. In this case, having smaller and fewer homoeologous rearrangements improved meiotic stability but not seed production as observed in ‘RCC’ and to a lesser extent in ‘CRC’. Although the number of new homoeologous rearrangements seemed to be limited as soon as the 6th generation, individuals still carry a large deleterious load on fertility and meiotic behavior that would not be purged in these self-fertilized lines. Hence, these fixed homoeologous rearrangements may only be eliminated from the population by extensive outcrossing events.

### 3. On the road to polyploid success

Homoeologous rearrangements are visible in *B. napus* varieties (Chalhoub *et al*., 2014; Lloyd *et al*., 2018; Song *et al*., 2020). These rearrangements are globally shorter in natural varieties compared to resynthesized lines (Chalhoub *et al*., 2014) and do not seem to influence meiotic behavior and fertility (Rousseau-Gueutin *et al*., 2017). Recently, the pangenome of *B. napus* revealed the high proportion of homoeologous genomic rearrangements in modifying important adaptive and agronomic traits, such as flowering time, seed quality, silique length disease resistance and chemical defense (Pires *et al*., 2004; Zhao *et al*., 2006; Hurgobin *et al*., 2017; Stein *et al*., 2017; Song *et al*., 2020). It is thus obvious that these rearrangements are occurring and may sometime be beneficial to agronomically improve *B. napus* varieties. Yet, these homoeologous rearrangements observed in natural *B. napus* never level up to the number and size of homoeologous rearrangements observed in newly resynthesized *B. napus* lines (Szadowski *et al*., 2010, 2011; Rousseau-Gueutin *et al*., 2017). Thus, we can hypothesize that the original *B. napus* population have experienced similar homoeolog rearrangements as observed in the resynthesized allotetraploids, but that overlapping between rearrangements of different sizes in the same genomic region allow crossover formation leading to progressive size decrease of homoeologous rearrangements. Thereby, the current *B. napus* varieties most presumably derive from a small founding population of euploid and fertile individuals that intercrossed to minimize the number of fixed homoeologous rearrangements, hence maximizing the number of bivalent and enhancing seed production. The alternative (or complementary) explanation lies in the presence of a genetic determinant that completely or partially prevented homoeologous pairing and rearrangements, and preexisted in *B. napus* diploid progenitors.

## CONCLUSION

To conclude, the consequences of homoeologous rearrangements on meiotic behavior and fertility depends on their size and the genetic background where they occur. Although recurrent euploid selection drastically reduced the fixation of novel homoeologous rearrangements in subsequent generations, it did not directly improve fertility and meiotic stability. Interestingly, the rearrangements identified in these *B. napus* enabled the identification of several candidate regions involved in seed yield and genome stability. These results offer a new perspective on the consequences of structural variants in allopolyploid genome stability and speciation success as well as new avenues to increase phenotypic diversity in oilseed rape.

## ACKNOWLEDGEMENTS

This work was made possible by financial support of the European Union and a Marie Sklodowska-Curie grant (number: 791908) awarded to JFC, as well as a grant from the department ‘Biology and Plant Breeding’ at INRAE (project ‘Allogen’ awarded to MRG). We acknowledge the BrACySol BRC (INRA Ploudaniel, France) that provided us with the seeds of parental lines; and K. C. Falk from Agriculture and Agri-Food Canada (Ontario) that provided the seeds of *B. rapa* ‘Z1’. We would also like to thank all the technical staff of the greenhouse for management of the plant material (especially L. Charlon, P. Rolland, J.-P. Constantin, J.-M. Lucas and F. Letertre). The authors have no conflict of interest to declare.

## AUTHOR CONTRIBUTION

AMC was involved in the experimental design with JFC and MRG in the conceptualization of the study; FE, ML, GT carried out the molecular and cytological experiments; MMG was in charge of plant care; JFC, SS, JM, CF, AMC, MRG were involved in data analyses; JFC, SS, AMC and MRG were involved in writing.

