## Supporting Information Fig. S1-S4 and Table S2 for "Untangling structural factors and evolutionary drivers in nascent polyploids"

The following Supporting Information is available for this article:

**Fig. S1** **Impact of artificial euploid selection.**

**Fig. S2** **Differences in fertility and meiotic behavior among resynthesized *B. napus* populations.**

**Fig. S3** **Evolution of structural rearrangements across all generations.**

**Fig. S4 Correlations between variables in all four resynthesized B. napus populations.**

**Table S1 Data table of phenotypic measures and structural rearrangements.**

**Table S2 Results of the Redundancy analysis.**

**Table S3 Results of the Genome scan analysis.**

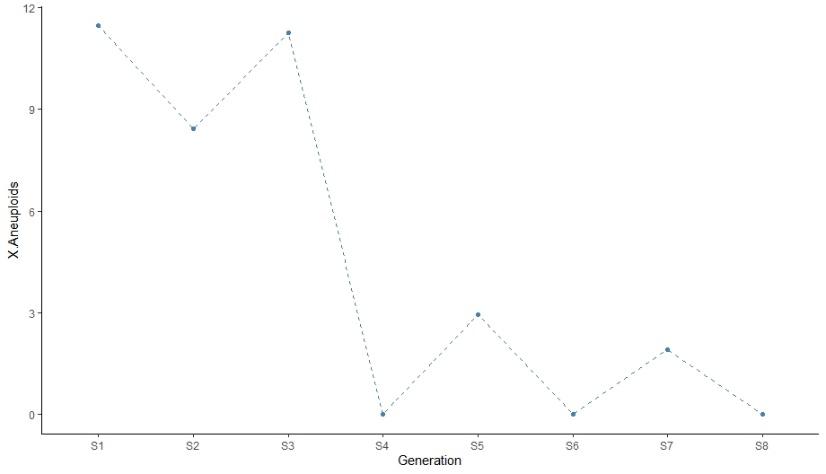
**Fig. S1 Impact of euploid selection on the percentage of aneuploid individuals at each generation overall resynthesized *B. napus* populations.**

**Fig. S2 Differences in fertility and meiotic behavior among resynthesized *B. napus* populations ‘RCC’, ‘CRC’, ‘EMZ’ and ‘UG EMZ’ (A) and *B. napus* var ‘Darmor’ for the number of seeds per 100 flowers; (B) for the percentage of cells with 19 bivalents and (C) for the percentage of cells with multivalents during Metaphasis I. Letters above graphs represent significative results of t-test (p<0.01).**

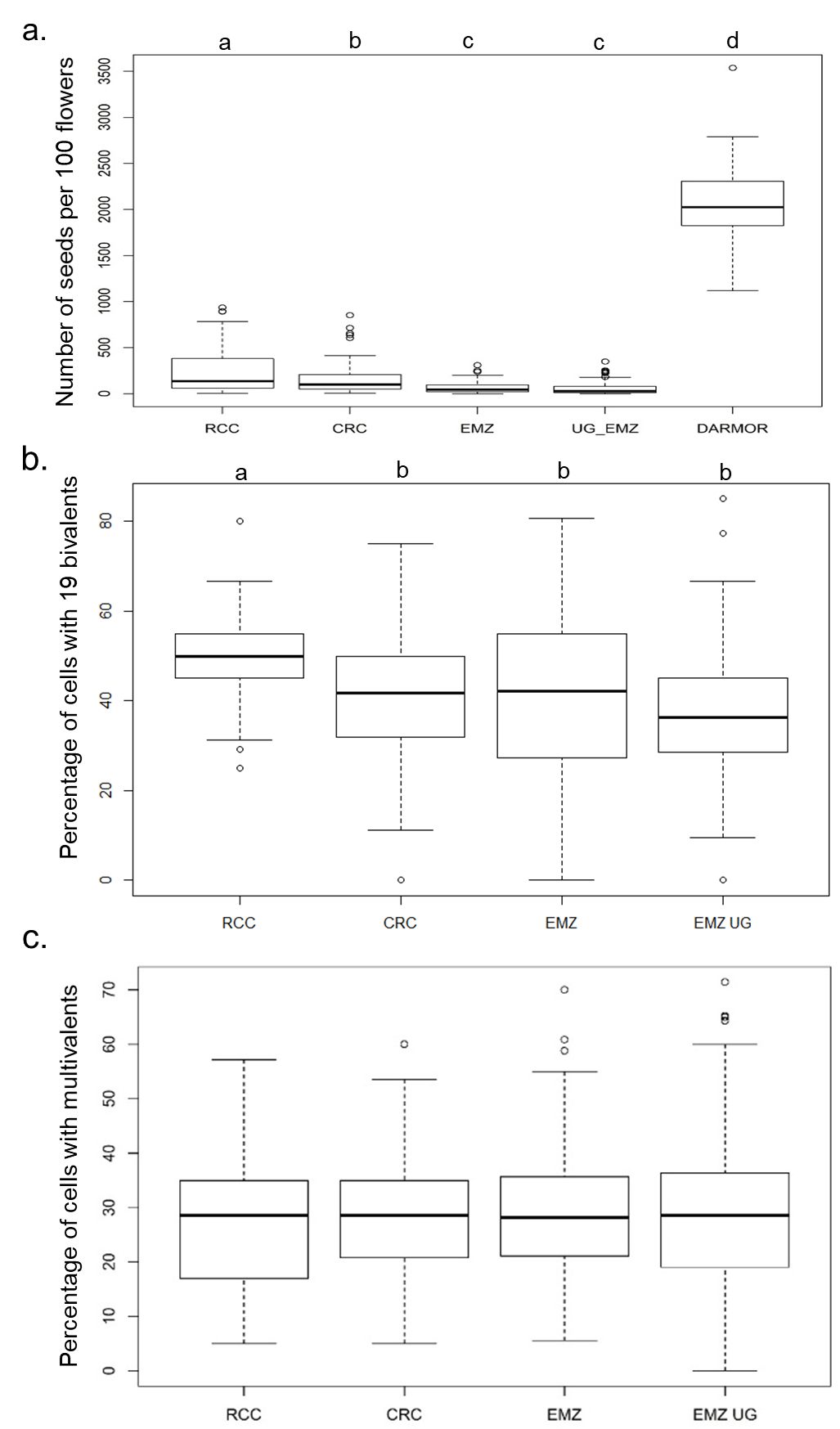

**Fig. S3 Identification of new, similar in size (Sim) or elongated (Ext) non-reciprocal fixed rearrangements at each generation for all four resynthesized *B. napus* populations.**

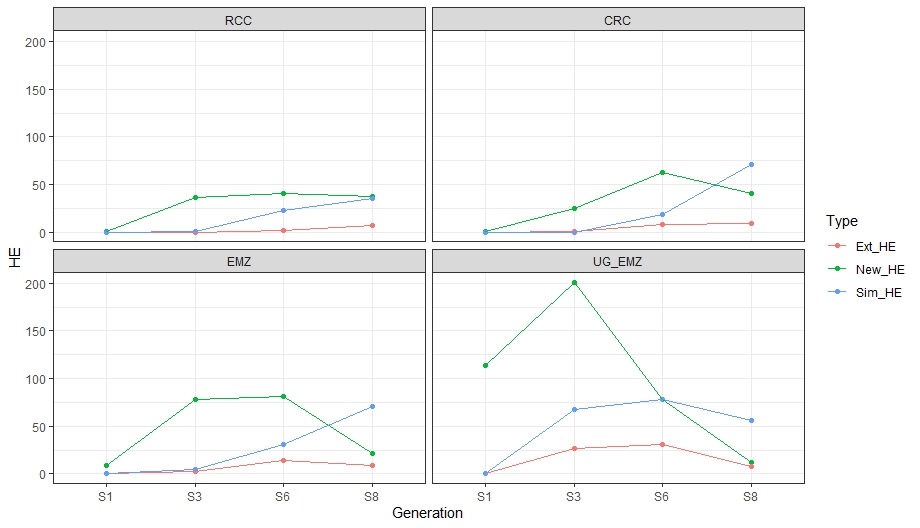

**Fig. S4 Pearson’s correlations were performed between the most explicative variables: percentage of cells with 19 bivalents, percentage of cells with multivalents, number of seeds per 100 flowers, survival at the next generation, number, average size and cumulative size of non-reciprocal homoeologous exchanges in (a) ‘RCC’, (b) ‘CRC’, (c) ‘EMZ’ and (d) ‘UG EMZ’. Color and size of circles are proportional to the correlation coefficients, only correlations with p<0.01 are presented with circles.**

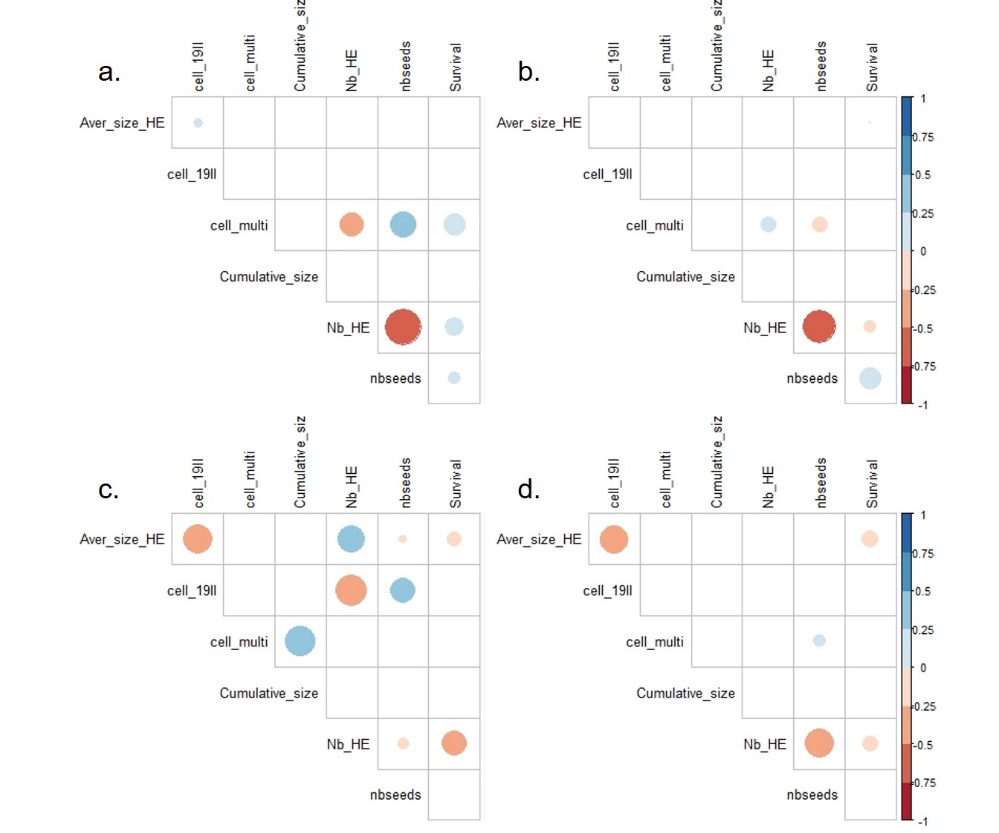

**Table S1** **Data table of phenotypic measures and structural rearrangements in resynthesized B. napus populations.**

SuppInfo_tableS1.xlsx

**Table S2** **Results of the Redundancy Analysis using the model: data_q ~ FG + Generation + FG:Generation, data = data_rda, which explains 68% of the variance at P-value < 0,001.**

|  | Df | Variance | F | Pr(>F) |
| --- | --- | --- | --- | --- |
| Population | 3 | 1.8628 | 23.4101 | 0.001*** |
| Generation | 4 | 5.7623 | 54.3129 | 0.001*** |
| Population*Generation | 12 | 1.3020 | 4.0907 | 0.001*** |
| Residual | 144 | 3.8194 |  |  |

**Table S3 Results of the Genome scan analysis.**

SuppInfo_TableS3.xlsx
